# Coexpression enables multi-study cellular trajectories of development and disease

**DOI:** 10.1101/719088

**Authors:** Brian Hie, Hyunghoon Cho, Bryan Bryson, Bonnie Berger

## Abstract

Single-cell transcriptomic studies of diverse and complex systems are becoming ubiquitous. Algorithms now attempt to integrate patterns across these studies by removing *all* study-specific information, without distinguishing unwanted technical bias from relevant biological variation. Integration remains difficult when capturing biological variation that is distributed *across* studies, as when combining disparate temporal snapshots into a panoramic, multi-study trajectory of cellular development. Here, we show that a fundamental analytic shift to gene *coexpression* within clusters of cells, rather than gene expression within individual cells, balances robustness to bias with preservation of meaningful inter-study differences. We leverage this insight in Trajectorama, an algorithm which we use to unify trajectories of neuronal development and hematopoiesis across studies that each profile separate developmental stages, a highly challenging task for existing methods. Trajectorama also reveals systems-level processes relevant to disease pathogenesis within the microglial response to myelin injury. Trajectorama benefits from efficiency and scalability, processing nearly one million cells in around an hour.

## Introduction

Single-cell RNA-sequencing (scRNA-seq) studies now profile millions of transcriptomes across diverse tissues, conditions, species, and ages^1–9^. To enable integration of biological patterns into multi-study insight, several algorithms have been developed to align common cell types across studies and then transform the underlying data to remove any study-specific differences^10–17^; cells deemed to be of the same cell type will thus have similar transcriptomic signatures in downstream analysis.

Unfortunately, because current integrative algorithms do not distinguish technical bias from real biological variation, they remove any meaningful change in a cell type across experimental conditions. A major task within single-cell analysis, however, is to infer trajectories and “pseudo-temporal” relationships among cells, thereby algorithmically reconstructing important continuous processes like differentiation or disease progression^18–21^. Reconstructing such trajectories across disparate studies, separated by both experimental bias and real cellular change, remains difficult even with state-of-the-art integration. Single-cell trajectories, therefore, remain practically limited to patterns observed within a single study.

Here, we unite both integration and trajectory inference, two major single-cell analytic efforts that have largely remained separate because current algorithms fail to achieve a delicate balance between robustness to unwanted bias and preservation of relevant multi-study variation (**Figure 1a**). To reveal dynamic biological processes at an unprecedented scope, we aim to construct *multi-study trajectories* of cellular change.

**Figure 1.**
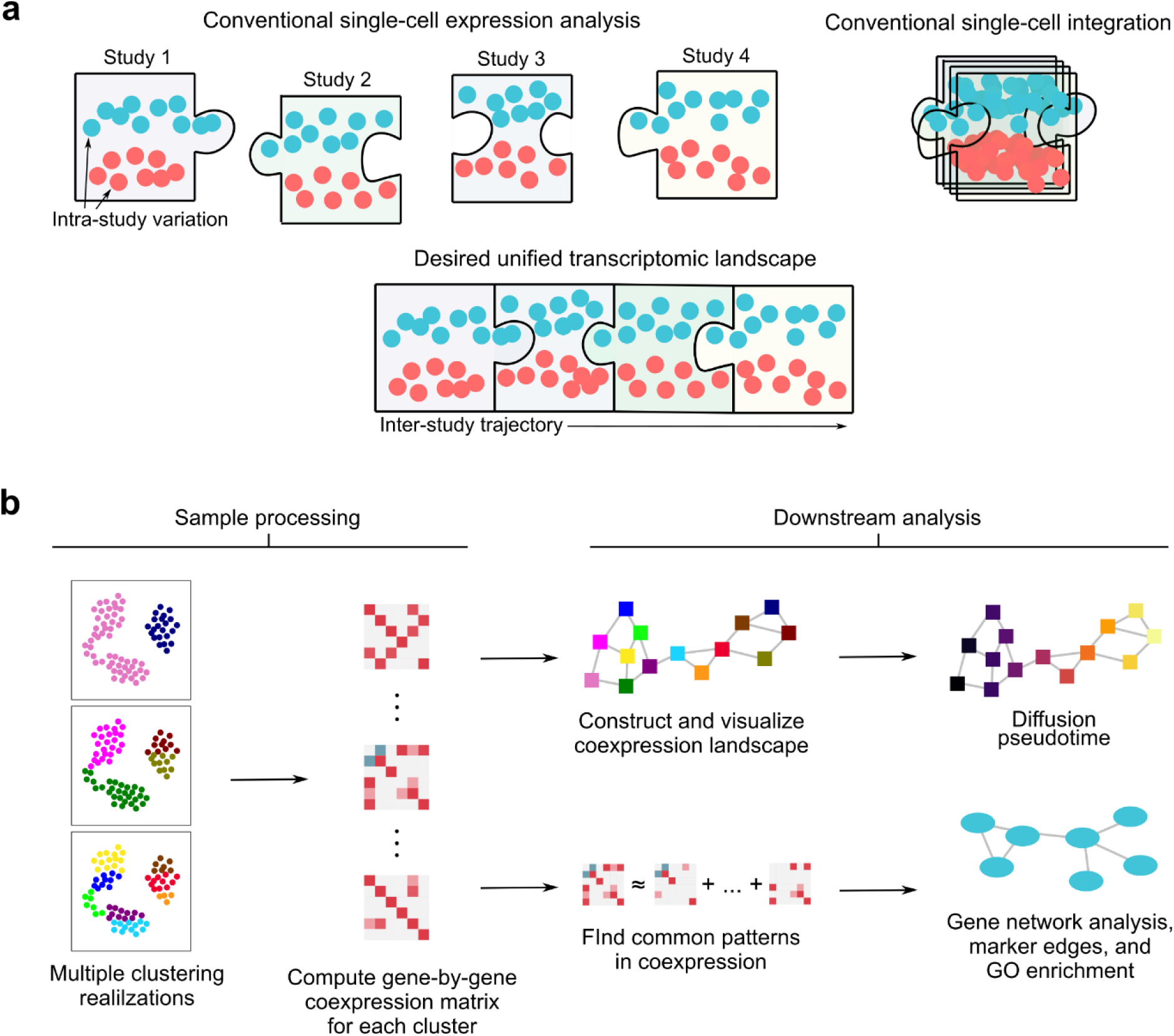
Overview of coexpression-based single-cell transcriptomic analysis. (**a**) A conceptual illustration of the difference between attempting to extract biological information from single-studies, each profiling different parts of a larger biological system (“Conventional single-cell expression analysis”); integrative algorithms that attempt to minimize inter-study variation but may also remove overarching biological structure, including temporal dynamics (“Conventional single-cell integration”); and piecing together structure across multiple studies of complex and dynamic biological systems, which we accomplish with single-cell coexpression (“Desired unified transcriptomic landscape”). (**b**) Overview of coexpression-based analysis, in which the fundamental analytic unit is a group of cells featurized by coexpression, rather than a single cell featurized by expression. Many downstream analyses have analogs in standard single-cell expression analyses.

Our novel, key insight is that differences in coexpression could preserve enough biological variation while still enabling integration. Coexpression is a conceptually favorable paradigm for integration since it favors redundant signal consistent across many genes^10,25–27^ and since common coexpression measures (e.g., Spearman correlation) are robust to many transformations of the data resulting from technical bias. In previous studies, coexpression has been used extensively to assess global gene expression changes in different biological conditions using both single-cell and bulk transcriptomics^22–24^; here, we show that analysis that respects variation in coexpression, combined with coexpression’s integrative properties, achieves a balance crucial to enabling multi-study trajectory inference.

We therefore introduce Trajectorama, a coexpression-based algorithm for integration that preserves and highlights cellular change across studies. Using Trajectorama, we efficiently integrate trajectories of neuronal development (across embryonic, neonatal, adolescent, and adult neurons) and hematopoiesis (across bone marrow, cord blood, fetal thymus, and peripheral blood) that no other integrative method is able to recover. Trajectorama’s coexpression feature space is highly interpretable, allowing us to probe the poorly understood microglial response to myelin injury, revealing a disease-associated gene network across demyelination models in mice and multiple sclerosis in human patients that implicates contributors to neurodegeneration.

Our conceptual advances beyond multi-study coexpression include panresolution clustering, in which we consider all clusters across a cellular hierarchy for downstream analysis, and interpretation through dictionary learning and functional analysis of condition-specific coexpression networks. Our algorithmic innovations and versatile applications—from understanding development across an entire lifespan to probing cell state change in response to disease—underscore the utility of coexpression-based trajectory integration.

## Results

### Multi-study coexpression analysis: Key concepts

In conventional single-cell transcriptomic analysis, the fundamental analytic unit is an individual cell described by features that encode levels of gene expression. A crucial difference in Trajectorama’s coexpression-based analysis is that the fundamental analytic unit is a *cluster* of cells; this cluster is in turn described by features that encode the *correlation* in expression between *pairs* of genes.

First, therefore, we require cells to be assigned to clusters. Clusters can be determined based on experimentally-determined properties or conditions, or such clusters can be determined by algorithms that group cells based on relative similarity in an unsupervised fashion^28^. While many clustering algorithms partition the data such that each cell is assigned to a single cluster, this need not be the case. Indeed, cells often belong to a hierarchy of biologically-meaningful groups^20^; for example, in brain tissue, it may be useful to separate neurons and glia, but within each category are many neuronal or glial subtypes. Rather than cluster cells based on a single level of a cellular hierarchy, i.e., a single clustering “resolution,” it is also possible to consider *all* clusters at *multiple* resolutions. This approach is particularly useful when determining clusters for coexpression-based analysis, since coexpression may change with clustering resolution^24,27^. We refer to this strategy as panresolution clustering, or *panclustering*.

After we determine clusters, each cluster is considered as a single datapoint in subsequent analysis. The features that describe a cluster are the correlations in expression (within that cluster) between all pairs of genes (**Figure 1b**). If there are *M* genes, then there will be 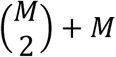 unique gene pairs, where we compute a correlation for each pair. In Trajectorama, we use the Spearman rank correlation due to its invariance under monotonic transformations of the underlying data and robustness to small numbers of large-magnitude outliers. Equivalently, we can think of each cluster as being described by a single gene-by-gene correlation matrix.

Equivalently, we can also think of each cluster as being described by a different gene association network, where the weights of edges connecting genes correspond to correlation strength. We can impose additional quality control cutoffs by setting low correlations to zero, or “sparsifying” the features, which helps reduce noise and improve computational efficiency, a property we take advantage of in our analysis.

Once we have featurized our clusters by coexpression, we can perform downstream analyses, many of which are analogous to standard expression-based analyses. For example, we can form trajectories by constructing a k-nearest-neighbors (KNN) graph where each node is a cluster and edges between nodes are added based on proximity in coexpression feature space. We can also find similarities and differences in coexpression among clusters, which correspond to stable or changing gene-gene associations. Correlations unique to a condition can in turn be interpreted as edges in a condition-specific gene network.

Trajectorama leverages and implements all of these concepts, encompassing cell clustering through coexpression featurization through downstream interpretation, within a single analytic framework, illustrated in **Figure 1b**. In particular, we design Trajectorama to integrate vast amounts of data while preserving relevant study-specific biological variation.

### Unified trajectory of neuronal development containing 932,301 cells

We first assessed whether coexpression could achieve the difficult balance of preserving continuously changing cellular phenotypes while overcoming study-specific bias. Given a wealth of scRNA-seq datasets that profile the mouse brain at different developmental timepoints, we reasoned that coexpression could construct a picture of neuronal development at an unprecedented scale. Known developmental age would help us validate the structure found by our analysis.

We therefore used Trajectorama to analyze five large-scale studies of mouse neurons from embryonic to adult. The first study^1^ used sci-RNA-seq3 to profile 562,272 cells representing the neural tube and notochord collected at day-length intervals from embryonic day (E)9.5 through E13.5. The second^3^ used Drop-seq and 10x Chromium v2 to profile 50,363 cortical neurons from late embryonic (E13.5 - E14.5) and postnatal day (P)10. The third^2^ used Microwell Seq to profile 10,796 cells across three developmental timepoints for E14.5, P1, and P56. The fourth^4^ used 10x Chromium v1 to profile 101,213 neurons from multiple adolescent timepoints from P12 through P27 and from a P60 adult. The fifth^5^ used Drop-seq to profile 207,657 neurons from P60 through P70 adults. This data was generated by laboratories spanning both United States coasts and three continents using single-cell or single-nucleus transcriptomic platforms and in total profiled more than 150 individual mice.

We obtained a panclustering of cells based on the Louvain community detection algorithm^29^, a common clustering method for scRNA-seq data. Louvain clustering iteratively merges cells into cluster “communities” until convergence, which is controlled by a resolution parameter^30^ (higher resolutions tend to increase the number of communities). We also obtain many possible realizations of a Louvain clustering by repeating the algorithm with multiple resolution parameters and use cluster assignments across all agglomerative iterations (**Methods**). To see if coexpression could directly overcome study-specific bias, we panclustered each study separately before combining clusters across all studies during downstream analysis in coexpression space.

When we visualize the coexpression landscape with a force-directed embedding^31^ of the KNN graph in which each node is a panresolution cluster, the graphical topology naturally arranges according to biological age (**Figure 2a**) rather than study-specific structure (**Figure 2b**). Analogous to assigning pseudotimes to cells in gene expression space, we can likewise run a diffusion-based pseudotime (DPT) algorithm^19^ within the coexpression landscape using the cluster with the lowest average age as the root of the diffusion process. Pseudotimes assigned to panresolution clusters in coexpression space were significantly correlated with biological age (Spearman *r* = 0.87, *P* < 10^−308^, *n* = 2,442 panresolution clusters) (**Figure 2c**).

**Figure 2.**
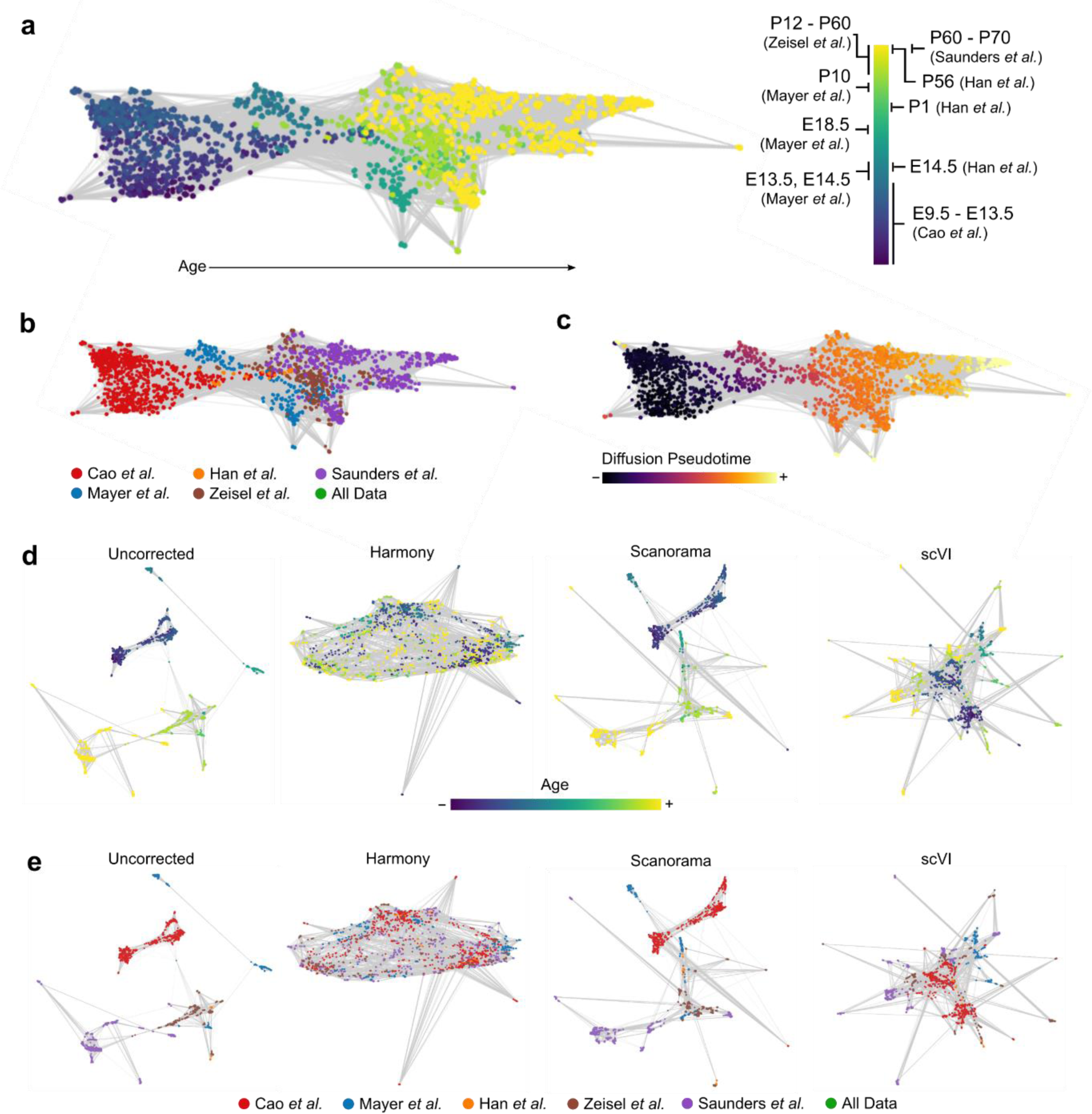
Coexpression landscape of mouse neuronal development. (**a**) A force-directed layout of the *k*-nearest-neighbors graph of panresolution clusters in coexpression space, which we refer to as the “coexpression landscape,” reveals a trajectory consistent with developmental age. (**b**) Studies are arranged according to order in developmental time, without removing all study-specific signal. (**c**) Diffusion pseudotime starting from the lowest-age node is strongly associated (Spearman *r* = 0.87, *P* < 10^−308^, *n* = 2,442 panresolution clusters) with biological age. (**d,e**) Panresolution clusters in uncorrected expression space and after correction with Scanorama or scVI still show large study-specific patterns without clear age-related structure. Harmony integration removes all study-specific differences including those related to developmental age.

If instead we use gene expression to learn two-dimensional visualizations of these datasets by plotting panresolution clusters using average gene expression, the datapoints arrange according to study-of-origin, without conveying any continuous developmental structure (**Figure 2d,e**). Uniform Manifold Approximation and Projection (UMAP) visualization of cells, the key algorithm underlying the Monocle 3 trajectory inference algorithm^1^, also does not convey the developmental relationships among the studies (**Supplementary Fig. 1**). Study-specific structure is still present after applying existing integrative algorithms based on mutual nearest neighbors matching^32^ (Scanorama) or on a latent space parameterized by a variational autoencoder (scVI)^16^ (**Fig. 2d,e**); these methods are representative of many others also based on nearest neighbors matching^11–13^ or on learning a joint latent space^10,15,17^. Another integrative method, Harmony^14^, removes nearly all study-specific signal, as designed (**Figure 2d,e**), which includes the valuable development-related information that only the coexpression landscape captures.

### Interpretation of coexpression landscape yields insight into neuronal development

Given this panoramic view into neuronal development, we facilitate further interpretation by highlighting similar coexpression patterns across many panresolution clusters with *dictionary learning*. In dictionary learning, we represent the coexpression matrix of each panresolution cluster as a sparse weighted sum of a few basis coexpression matrices, or “dictionary entries.” Each basis matrix can also be interpreted as a network, with edges between genes weighted by coexpression. Dictionary learning for correlation matrices has been successfully applied to diverse problems, including information retrieval^33^ and functional brain profiling^34^.

We looked for significant gene ontology (GO) process enrichments^35^ within the set of genes involved in “marker edges” unique to a particular dictionary entry, using a background set of all genes considered in our coexpression analysis (around two thousand highly variable genes; **Methods**). Within the embryonic portion of the coexpression landscape, we observe differentiation and developmental processes (GO:0051094, false discovery rate [FDR] *q* = 3.3 × 10^−3^) and neuron fate commitment (GO:0048663, FDR *q* = 8.4 × 10^−3^). Late-fetal and early-postnatal development includes neurogenesis (GO:0050767, FDR *q* = 3.9 × 10^−4^) and neuron projection organization (GO:0030030, FDR *q* = 0.018). Adolescent and adult stages are enriched for a more diverse set of processes from neurotransmission (GO:0001505, FDR *q* = 1.5 × 10^−4^) to amyloid-β response (GO:1904646, FDR *q* = 0.042). The enriched processes for all of these dictionary entries are consistent with their respective developmental stages, offering evidence that Trajectorama integration preserves inter-study patterns due to biological development.

We can also look at individual genes that are strongly associated with diffusion pseudotime in the coexpression landscape and validate them with the Allen Developing Mouse Brain Atlas (ADMBA), which spatially locates the expression of around 2000 preselected genes using in situ hybridization (ISH) experiments^36^. Genes with the strongest associations with developmental pseudotime also showed strong developmental changes in ISH intensity in the expected direction, i.e., increasing or decreasing with development. The top such positively correlated gene is *Fos* (Spearman *r* = 0.67; *n* = 2,442 panresolution clusters), which encodes a well-known marker of neuronal activity^37^; the top such negatively correlated gene is *Eomes* (Spearman *r* = −0.45; *n* = 2,442 panresolution clusters), which encodes an important transcription factor in early neurogenesis^38^ (**Figure 3b,c**).

**Figure 3.**
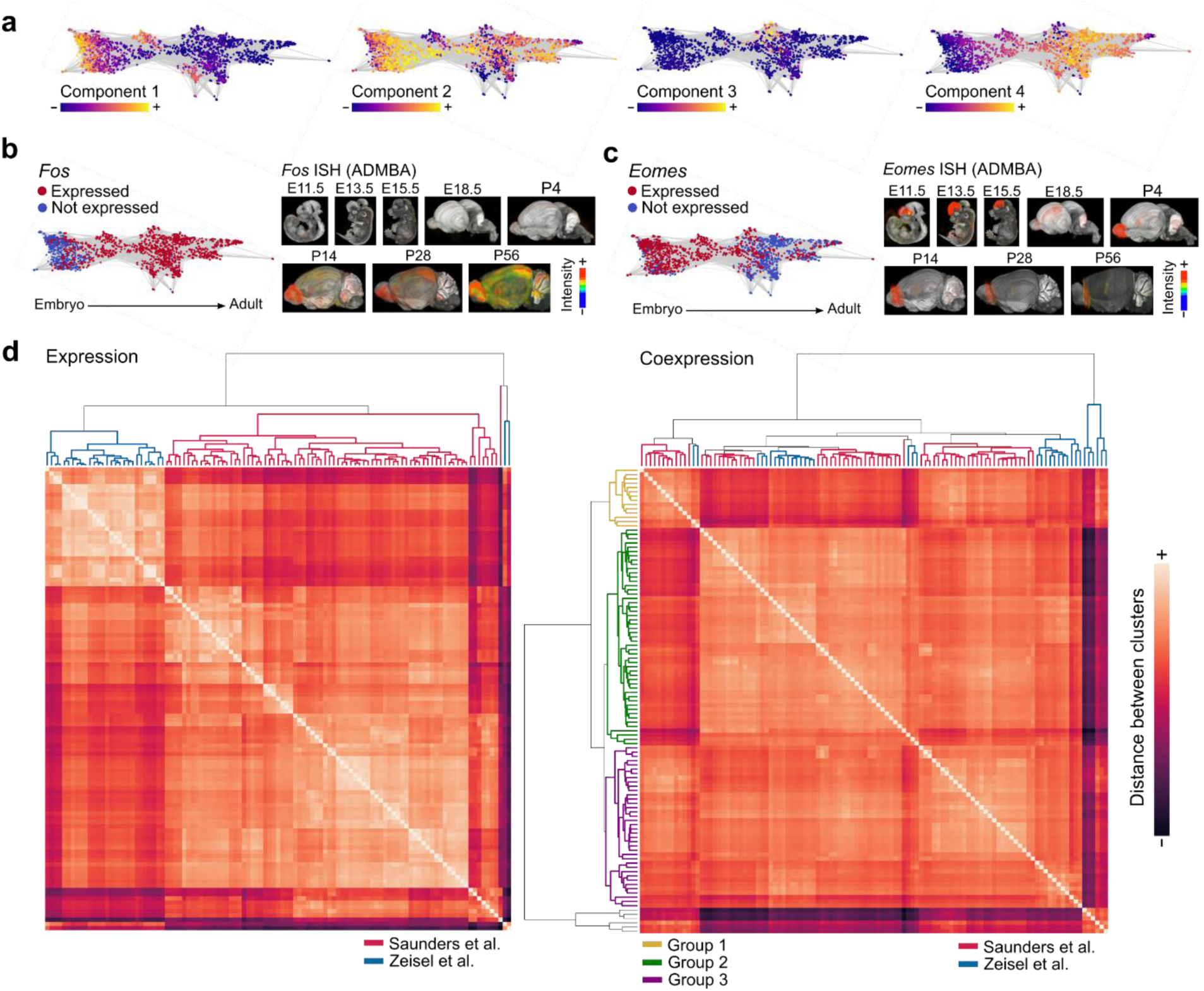
Neuronal trajectory interpretation and cross-study subtype integration. (**a**) Dictionary entries highlight different stages of neuronal development. (**b,c**) We observe positive correlations between diffusion pseudotime, corresponding to development, with the expression of genes such as *Fos* and negative correlation with the expression of *Eomes*. Changes in expression of these genes over development are validated and spatially located by the Allen Developing Mouse Brain Atlas (ADMBA)^36^. Images show locations and levels of gene expression intensity measured by in situ hybridization (ISH); blue-green is low, yellow-orange is medium, and red is high. (**d**) Neuronal subtypes featurized by mean expression group primarily according to study while subtypes featurized by coexpression group primarily according to three main groups, followed secondarily by study.

Our analysis also reveals genes strongly associated with development, such as *Gm9945* and *Pon3* (Spearman *r* = 0.78 and *r =* −0.54, respectively; *n* = 2,442 panresolution clusters), that the ADMBA did not include in their list of assayed genes but may be important to include in future developmental studies. We make these correlations and GO enrichments available as **Supplementary Data**, which may be of further interest to developmental biologists.

### Neuronal developmental landscape is robust to parameter choice

Two important parameters control the amount of information considered in our analysis and can be thought of as “smoothing” parameters. The first is the correlation cutoff parameter that controls the sparsity of underlying correlation matrices; lower values include more information but may increase noise and computational burden. The second is the number of nearest neighbors in the KNN graph representing the coexpression landscape, which impacts both visualization and diffusion pseudotime; considering more nearest neighbors results in a smoother trajectory. While we do introduce some smoothing into our analysis, the studies are consistently arranged according to their developmental order even as these parameters vary. With less smoothing, we also observe age-related branching of the developmental trajectory, suggestive of neuronal subtype-related structure (**Supplementary Fig. 2**).

### Coexpression integrates neuronal subtypes across studies

While the most pronounced signal captured within the neuronal trajectory is developmental age, there is still substantial heterogeneity among neurons. We therefore sought to determine if Trajectorama could provide multi-study insight into neuronal *subtypes* as well. To do so, we relied on extensive expert labelling of neuronal subtypes from Zeisel *et al*.^4^ (adolescent mice) and Saunders *et al*.^5^ (adult mice) to define neuronal clusters of interest. When comparing these subtypes in gene expression space, subtypes group primarily according to study (**Figure 3d**). When we instead featurize by coexpression, the clusters group primarily according to common subtypes, and only secondarily (since we do expect some differences due to real biological change) according to study (**Figure 3d**).

Neuronal subtypes group according to three major coexpression-based patterns. Genes most unique to the first group are enriched in glutamergic structures (GO:0098978, FDR *q* = 1.5 × 10^−11^) and glutamate signaling (GO:0035235, FDR *q* = 6.4 × 10^−3^). In contrast, the second group has significant enrichments for both adrenergic (GO:0004935, FDR = 0.017) and cholinergic (GO:0032224, FDR *q* = 0.037) processes. The third group, which also contains the highest number of adolescent subtypes, is most significantly enriched for neurons with synaptic plasticity (GO:0048167, FDR *q* = 9.3 × 10^−8^) and involved in cognition (GO:0007611, FDR *q* = 1.9 × 10^−4^), learning, and memory (GO:0007611, FDR *q* = 7.4 × 10^−5^). The hierarchy of subtypes has additional structure as well, though we focused on only the three largest, highest-level groupings that each contain subtypes from both studies (**Figure 3d**).

### Trajectorama constructs a multi-tissue hematopoietic trajectory

Based on the ability of Trajectorama to integrate neuronal studies while respecting biological change, we next set out to establish if it could demonstrate similar capabilities within a completely separate developmental system. To this end, we analyzed the coexpression landscape of four hematopoietic datasets from the fetal thymus^39^, bone marrow, cord blood^7^, and peripheral blood^6^. Throughout these tissues, we expect to observe cells in many stages of hematopoiesis, including stem cells and erythroid progenitors, mostly in the bone marrow and cord blood, to more mature lymphocytes and myeloid cells, mostly as peripheral blood mononuclear cells (PBMCs)^40^.

Visualizing the coexpression landscape of panresolution clusters obtained across all studies reveals an organization consistent with the three main branches of hematopoiesis corresponding to erythropoiesis, myelopoiesis, and lymphopoiesis (**Figure 4a**). Such organization (with similar developmental granularity) has been observed in the gene expression space^20^ and in the chromatin accessibility space^41^ of single studies in single tissues, but, importantly, here we instead show a unified hematopoietic landscape across multiple tissues generated by disparate laboratories.

**Figure 4.**
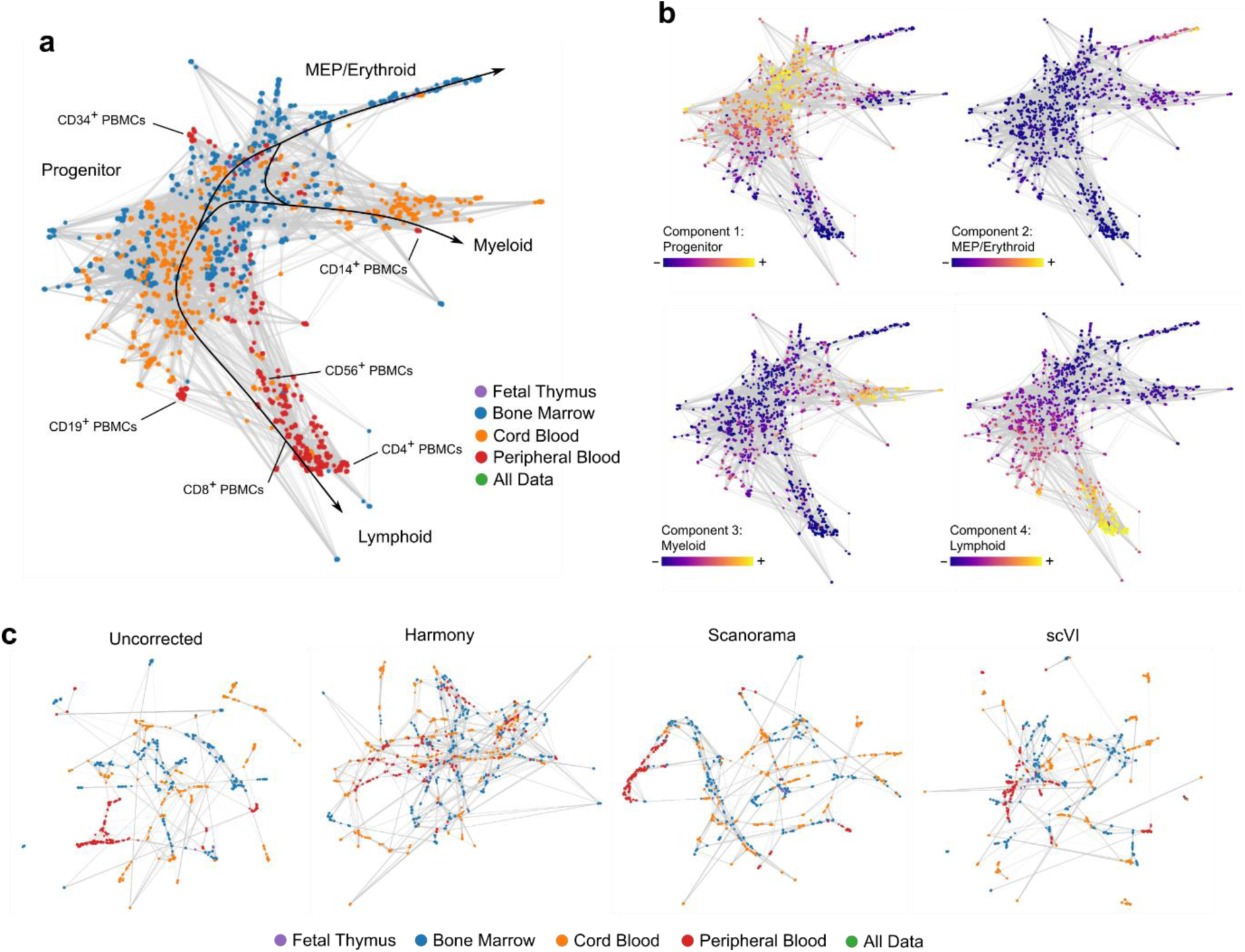
Coexpression landscape of human hematopoiesis. (**a**) The coexpression landscape of immune cells from bone marrow, cord blood, and peripheral blood organizes largely according to erythropoietic, lymphopoietic, and myelopoietic lineages. Some of the PBMCs have FACS-derived labels, enabling us to place clusters with known surface markers in various regions of the coexpression landscape (also see **Supplementary Fig. 3**). (**b**) Dictionary learning of the coexpression matrices separates the coexpression landscape into four main regions; FACs labels and GO process enrichments suggests that these dictionary entries correspond to the different, main stages of hematopoiesis. (**c**) Existing integrative methods either do not overcome study specific bias (Scanorama and scVI) or obscure the lineage relationships among the four tissues (Harmony).

We interpret different branches in the coexpression trajectory partially based on experimentally-determined PBMC labels. Prior to scRNA-seq, a large number of the PBMCs underwent fluorescence activated cell sorting (FACS) for progenitor-associated (CD34^+^), myeloid-associated (CD14^+^), and lymphoid-associated (CD4^+^, CD8^+^, CD19^+^, CD56^+^) cell-surface marker expression (**Supplementary Fig. 3**). Dictionary learning yielded four main dictionary entries corresponding to the major regions within the landscape (**Figure 4b**). The first dictionary entry, which we call progenitor-associated, corresponds to all of the CD34^+^-labeled clusters. The second dictionary entry, which we call erythropoietic, includes GO enrichments related to heme biosynthesis (GO:0006783, FDR *q* = 0.04) and strong metabolic signatures (GO:0044237, FDR *q* = 1.0 × 10^−12^). The third dictionary entry, which we call lymphopoietic, includes all lymphoid-specific (CD4^+^, CD8^+^, CD19^+^, CD56^+^) clusters. The fourth dictionary entry, which we call myelopoietic, includes some CD14^+^ clusters and GO enrichments involving myeloid differentiation (GO:0045637, FDR *q* = 3.4 × 10^−4^).

We also note that PBMCs largely exist at the periphery of the landscape, consistent with such cells being the most mature within the hematopoietic lineage. In contrast, Harmony-based integration removes all tissue-specific differences and obscures the lineage relationships among the tissues (**Figure 4c**) while mean expression of clusters without correction, and even following Scanorama and scVI correction, primarily exhibits study-specific structure (**Figure 4c**). Overall, our hematopoietic analysis adds additional support for coexpression as an integrative strategy that can preserve key biological differences among disparate studies.

### Trajectorama reveals a disease-specific microglial gene network

While Trajectorama can yield panoramic views across long developmental scales, we next wanted to assess if it could also reveal more fine-grained insight into biological systems that are less well understood. In particular, recent work has begun to illuminate the key role of microglia in neurodegenerative disease^42,43^, for which coexpression provides a unique opportunity to integrate information across multiple microglial studies while still preserving disease-specific signal.

We therefore integrated microglia from mouse and human samples across three studies^5,8,9^, which together contained single-cell microglial transcriptomes from multiple points along a mouse lifespan, from models of mouse brain injury (facial nerve axotomy and demyelination), and from human donors with and without multiple sclerosis (MS). The Trajectorama coexpression landscape includes a main age-related trajectory, from embryonic (E14.5) through aged (P540) microglia, and off-trajectory outlier clusters from injured tissue samples (**Figure 5a**); similarly, hierarchically grouping the known microglial conditions based on similarity in coexpression space (**Methods**) obtains a clear outlier group consisting of microglia from mice that had undergone artificial demyelination and from human MS patients (**Figure 5b**). We note that, in coexpression space, this injury-associated group naturally separates from other microglial conditions without supervision.

**Figure 5.**
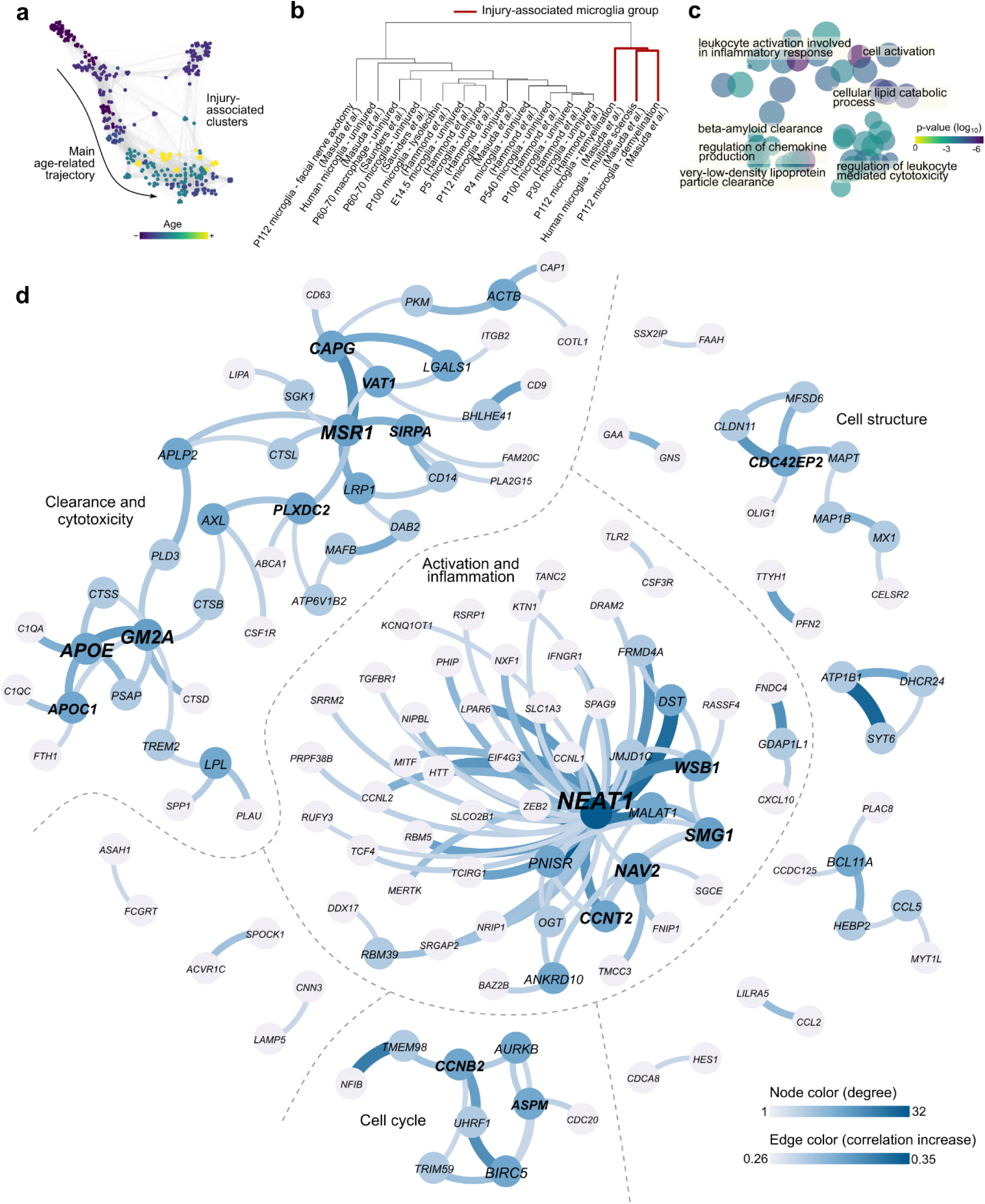
Multi-study analysis of microglial response to myelin injury. (**a**) The coexpression landscape of panresolution clusters reveals a main age-related trajectory as well as off-trajectory outlier clusters from injured tissue. (**b**) Grouping microglial subtypes reveals a cluster containing injury-associated conditions in both mouse and human microglia. (**c**) GO enrichment terms are visualized in two dimensional “semantic space” with key terms relevant to disease-associated microglia also displayed. (**d**) The disease-specific coexpression network reveals functional gene modules related to myelin injury. The top 150 edges in which coexpression increases from a baseline microglial state are arranged into a disease-associated coexpression network; almost all of these associations have not been described by previous studies. Major subgraphs are labeled according to GO terms associated with internal genes. Nodes are colored darker blue with higher degree; gene labels are larger and bolder with higher degree; and edges are thicker and darker blue with a higher increase in correlation from the baseline microglial state.

We then constructed an injury-associated coexpression network by considering the gene pairs with the highest increase in coexpression, combined across both mouse and human injury conditions, relative to baseline microglial coexpression (**Methods**). GO enrichment analysis of genes ranked by increased coexpression in disease state reveals three main functional categories: lipid and protein clearance, leukocyte-mediated cytotoxicity, and cellular activation involved in inflammation (**Figure 5c; Supplementary Data**). These processes are consistent with the hypothesized role of microglia in MS as involved in clearance of damaged myelin via phagocytosis^42^ and as drivers of neurodegenerative pathogenesis by inducing neuronal cell death^44^ and promoting local inflammation^42^.

The most valuable insight into microglial processes relevant to disease, and to myelin injury in particular, comes from visualizing the injury-associated coexpression network itself (**Figure 5d**). Two major connected components appear in the network: the first related to lipid clearance and leukocyte-mediated cytotoxicity and the second related to inflammatory activation.

The first connected component recovers key gene modules that have been implicated in neurodegeneration. Of special note, the network recovers an *APOE/TREM2/GM2A* gene module that has been extensively linked to a microglial “sensor” of neurodegeneration^9,43,45^. Another high-degree module includes *SIRPA*, which regulates demyelination repair^46^, and *MSR1*, which has been implicated in myelin uptake in MS lesions^47^. The network suggests a correlative link from the *APOE*/*TREM2* neurodegenerative sensing module to the *SIRPA*/*MSR1* uptake and clearance module through genes like *AXL*, which has also been suggested as essential to recovery from myelin injury^48^. While many of these genes have been *individually* implicated in neurodegeneration, we note that our coexpression-based analysis suggests *links* among these key genes that are useful for follow up study. Experimentally establishing the causal role these genes play in disease pathogenesis is important future work.

The second major connected component centers on the *NEAT1* long noncoding (lnc)RNA, which recently has been linked to inflammatory activation of macrophages^49^. These results suggest that the observation of *NEAT1* in MS serum, for which the mechanism was previously unknown^50^, is tied in part to microglial inflammation. Further experimentation is needed to see if *NEAT1* leads to or is a consequence of inflammation-mediated pathogenesis, or it could also serve as a biomarker of MS disease or related inflammation.

More broadly, the injury-associated microglia network illustrates how coexpression-based analysis across multiple experiments can generate further hypotheses that lead to novel biological discovery. Not only can coexpression analysis elucidate broad developmental changes, its rich feature space and inherent interpretability can also provide deep insight into cell state changes such as those in health versus disease.

### Trajectorama is practical for datasets with millions of cells

To enable consortium-scale analysis, we made algorithmic choices that allow scalability to large numbers of cells, while preserving the ability to model complex phenomena. For example, we choose to sparsify our coexpression matrices using a nominal cutoff rather than the memory intensive strategy of preserving dense correlation matrices or the runtime intensive strategy of learning sparse covariance matrices via regularization^51^ (**Supplementary Table 1**). Since scRNA-seq experiments typically measure little to no signal for many genes, we also limited analysis to around two thousand genes with highest statistical variability, a common dimensionality reduction strategy in conventional expression analysis^28,52^ (**Methods**).

We performed all of our analyses in a practical amount of computational time and resources. Our entire coexpression-based procedure, which includes panresolution clustering through downstream analysis of the coexpression landscape, analyzes almost a million cells in a little over an hour on a standard cloud instance with 16 cores (**Supplementary Table 2**). Our pipeline has a runtime and memory usage with a close-to-linear asymptotic scaling in the number of cells and a worst-case quadratic asymptotic scaling in the number of features (i.e., genes). While the coexpression space may seem cumbersomely quadratic, scRNA-seq experiments typically measure only around one or two thousand genes with nontrivial variability^52^; moreover, the number of strong correlations is usually within the same order of magnitude as the number of highly variable genes.

Once the data has been summarized as panresolution clusters, further downstream analysis including visualization, pseudotime assignment, and dictionary learning becomes extremely efficient due to the greatly reduced number of datapoints; in the case of mouse neuronal development, analysis is done on just 2,442 panresolution clusters instead of 932,301 single cells. The resource requirements for different stages of our analytic pipeline on the mouse neuronal development analysis are provided in **Supplementary Table 2**.

## Discussion

Our work shows that researchers can analyze an unprecedented amount of information across scRNA-seq studies, while retaining key biological variation, by focusing on the coexpression matrix of a group of cells as the fundamental unit of analysis. While not intended as a complete replacement for current integrative methods, as we have shown, Trajectorama can be valuable when researchers wish to integrate data while preserving inter-study biological variation. As laboratories continue to conduct single-cell experiments that explore heterogeneous biological models and conditions, we expect such scenarios to be ubiquitous.

By leveraging coexpression, Trajectorama benefits from a number of additional properties. Current integrative methods map cells into an arbitrary feature space that only preserves *relative* meaning (for example, cell A is more similar to cell B than to cell C). In contrast, coexpression has *intrinsic* meaning: each feature in coexpression space is simply the correlation between two genes (for example, Spearman correlation^53^), a fundamental and intuitive data science concept. Trajectorama is also highly efficient, since it combines information across many cells similar to existing algorithms that accelerate workflows via data sketching or summarization^54,55^.

Our results suggest many directions for future work. Our coexpression matrices are not positive semidefinite (PSD) for practical reasons, but efficiently learning large numbers of nontrivially sparse PSD matrices is an interesting and challenging task. If all coexpression matrices are PSD, it may be possible to leverage the distance along the manifold represented by all PSD matrices to obtain more natural dictionary learning-based decompositions^33^ and nearest-neighbor queries (which would involve designing new techniques for efficient nearest-neighbor search). Additional methods might also enforce further constraints within the dictionary learning objective (for example, basis matrices that are valid correlation matrices) or take other approaches to interpreting large numbers of coexpression matrices like common principal components analysis^56^ or other kinds of tensor decomposition^57^.

Other considerations include exploring alternative methods for measuring coexpression^58^, inferring causal gene regulatory networks, or exploring different clustering strategies, panresolution or otherwise. A larger question is whether other feature spaces exist that enable multi-study trajectories; for example, metric learning approaches could directly construct such a space via known developmental metadata^59^. Reasoning about the relationship between coexpression and other functional associations within single cells, like those involving chromatin accessibility or methylation, remains an important consideration.

Trajectorama can be used to probe biological systems beyond those interrogated in this study, providing an informative analysis that is complementary to existing integrative methods for studying biological processes at single-cell resolution and at multi-institution scale. We make our analysis pipelines and data available at http://trajectorama.csail.mit.edu.

## Methods

### Mouse neuronal development dataset preprocessing

We obtained publicly available datasets from five large-scale, published single-cell transcriptomic studies of the mouse brain at different developmental timepoints^1–5^. We used only the cells that passed the filtering steps of each respective study and additionally removed low-complexity or quiescent cells with less than 500 unique genes. For the embryonic dataset from Cao *et al*.^1^, we only considered cells that the study authors had assigned to the “neural tube and notochord” trajectory. For the datasets from Zeisel *et al*.^4^ and Saunders *et al*.^5^ we only considered cells that the study authors had labeled as neuronal. We then intersected the genes with the highest variance-to-mean ratio (i.e., dispersion) within each study to obtain a total of around 2,000 genes that were highly variable across all studies. All studies provided data as digital gene expression (DGE) counts, which we further log transform after adding a pseudo-count of 1.

### Human hematopoiesis dataset preprocessing

We obtained publicly available datasets of cord blood and bone marrow cells from the Human Cell Atlas^7^ (https://preview.data.humancellatlas.org/) and PBMCs from Zheng *et al*.^6^ (https://support.10xgenomics.com/single-cell-gene-expression/datasets). We removed cells with less than 500 unique genes; we also noticed a large number of cells with high percentages of ribosomal transcripts, which may indicate nontrivial amounts of ambient ribosomal RNA contamination during the scRNA-seq experiment, so we only included cells with less than 50% ribosomal transcripts in further analysis. As described previously, we intersected the genes with the highest dispersions within each study to obtain a total of around 2,000 genes that were highly variable across all studies. All studies provided data as digital gene expression (DGE) counts, which we further log transform after adding a pseudo-count of 1.

### Microglia dataset preprocessing

We obtained publicly available datasets from three single-cell transcriptomic studies of microglia across a diverse set of conditions^5,8,9^. We kept only the cells labeled by the original studies as microglia and we additionally removed low-complexity or quiescent cells with less than 500 unique genes. Mouse genes were mapped to human orthologs. As described previously, we intersected the genes with highest dispersions within each study to obtain around 2,000 genes that were highly variable across studies, followed by a log transformation after adding a pseudo-count of 1.

### Panresolution clustering

We modify the Louvain clustering algorithm^29,30^ (https://github.com/vtraag/louvain-igraph) to store community information at each iteration. We choose Louvain clustering due to its asymptotic efficiency, since its runtime and space usage scales with the size of the *k*-nearest neighbor (KNN) graph of cells (i.e., each cell is a node in the graph), rather than quadratically in the number of cells as in other hierarchical clustering algorithms. To capture a range of potential clustering results, we rerun the Louvain clustering algorithm at a diverse range of clustering resolutions (0.1, 1, and 10) on a 15-nearest neighbor graph, constructed using Euclidean distances in gene expression space, storing the hierarchical cluster information for each run. The three runs of Louvain clustering are done in parallel and we cluster each study individually. To reduce the effect of noisy correlations, we consider clusters with a minimum of 500 cells, which, combined with highly variable gene filtering (described below), reduces the chance that a strong correlation is due to a few outlier cells.

### Computing coexpression matrices

We compute the Spearman correlation matrix **R**^(*i*)^ ∈ [−1,1]^*M*×*M*^ for each of the panresolution clusters obtained as described above, where *i* ∈ {1,2, …, *N*} with *N* denoting the number of panresolution clusters and *M* denoting the number of highly variable genes. The entry 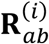 at row *a* and column *b* of **R**^(*i*)^, corresponding to the *a*^th^ and *b*^th^ genes, takes the value

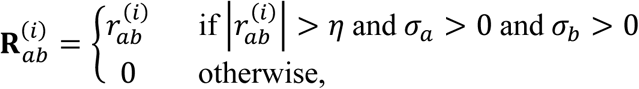

where 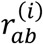 is the Spearman correlation coefficient^53^ and *σ*_*a*_ and *σ*_*b*_ are the respective standard deviations of the rank values of the gene expressions (which appear in the denominator of the Spearman correlation expression). *η* ∈ [0, 1] is a sparsification parameter that sets low correlations to zero and can be interpreted as a smoothing parameter that preserves only the most important associations. Low values of this parameter can introduce additional structure into the analysis, but may also introduce larger amounts of noise (see **Supplementary Fig. 2**).

### Visualization and diffusion pseudotime analysis of panresolution clusters

To visualize the coexpression landscape defined by the panresolution clusters, the symmetric correlation matrices **R**^(*i*)^ ∈ [−1,1]^*M*×*M*^ are treated as vectors 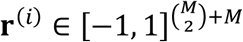 on which we construct the *k*-nearest neighbors graph using the Euclidean distance in coexpression space as the distance metric. This graph was visualized with a force-directed embedding using the ForceAtlas2 algorithm^31^ (https://github.com/bhargavchippada/forceatlas2). For the mouse neuronal development analysis, a diffusion pseudotime (DPT) algorithm^19^ was applied to this graph using the panresolution cluster with the earlies average age as the root. Larger values of *k* can also increase the amount of smoothing in the structure captured by the *k*-nearest-neighbors graph and subsequent visualization and DPT analysis (see **Supplementary Fig. 2**). We used the implementation in Scanpy^60^ (https://scanpy.readthedocs.io/en/stable/) for the *k*-nearest neighbors graph construction and DPT analysis.

We also visualized panresolution clusters in gene expression space, Harmony-integrated expression space^14^, Scanorama-corrected expression space^32^, and scVI-integrated latent space^16^. To summarize features across multiple cells into a single feature vector for each panresolution cluster, we used the mean expression. We similarly constructed the *k*-nearest-neighbors graph with panresolution clusters as nodes and Euclidean distance between the summarized gene expression values as the distance metric.

### Coexpression matrix dictionary learning

We formulated the dictionary learning problem for coexpression matrices by optimizing

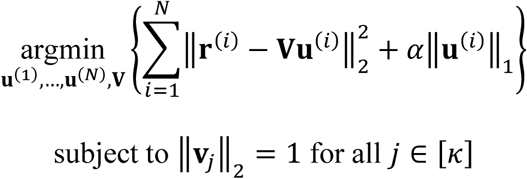

where 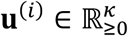 is a sparse code of weights for panresolution cluster *i, α* is a sparsity-controlling parameter, 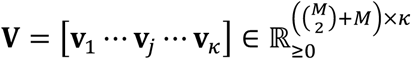 is a dictionary of *κ* (vectorized) coexpression matrices, and *κ* is a user-defined parameter indicating the number of dictionary entries to learn. We used an iterative optimization algorithm that alternatively estimated dictionary weights and dictionary entries using least angle regression-based optmiziation^61^ until convergence. We tune *κ* by plotting the objective function error versus values of *κ* and manually selecting a value after which there are relatively smaller drops in objective function values, a parameter selection procedure often referred to as the “elbow method.”

### Interpretation of dictionary entries

We can interpret each dictionary entry **v**_***j***_ as a coexpression network in which genes are nodes and elements of **v**_***j***_ define edge weights between those genes. We use the networkx Python package^62^ to represent graphs and compute various graph statistics. Using genes that are involved in edges that are unique to a given coexpression network, we look for gene ontology (GO) process enrichments using a background set of all highly variable genes considered in the analysis, for which *P*-values can be computed using a hypergeometric null model followed by subsequent FDR *q-*value computation^63^. We use the GOrilla webtool (http://cbl-gorilla.cs.technion.ac.il/)^35^ with default parameters, which reports all enrichments more significant than a nominal *P*-value of 1e-3. We use the REVIGO webtool (http://revigo.irb.hr/) with default parameters, which consolidates similar GO terms and visualizes terms in a two-dimensional “semantic space” that places similar terms closer together^64^. We only consider dictionary entries that have nonzero weights in at least ten panresolution clusters.

### Neuronal subtype hierarchical grouping and interpretation

Cellular subtypes were determined according to expert curated labels provided by the original studies^4,5^. Each subtype was featurized by coexpression and by mean expression for benchmarking purposes. Agglomerative hierarchical clustering of the subtypes was then performed using the scipy Python library^65^. To interpret genes unique to a group of subtypes, we computed the mean coexpression within the group and sorted each dimension according to the highest increase in correlation from the mean coexpression of all subtypes. Genes were then ranked according to the first appearance of the gene within the sorted list of coexpression dimensions; this gene ranking was used as input into the GOrilla webtool for GO enrichment analysis.

### Microglial subtype analysis and interpretation

Microglial subtypes were determined based on unique combinations of age, species, and tissue injury status. For the coexpression landscape analysis, each of these subtypes was considered as a separate study. Fewer clusters enabled a lower sparsification threshold of *η* = 0.1. All other methods and parameters remained the same.

As in the neuronal subtype analysis, we also hierarchically clustered the microglial subtypes and observed an injury-associated group of microglial subtypes. We took the coexpression mean of this injury-associated microglial group, including both mouse and human clusters, and compared it to the mean of all microglial subtypes. Coexpression dimensions were sorted according to the highest increase in correlation within the injury-associated group. This sorted list was used to rank genes as input into the GOrilla webtool for GO enrichment analysis and the first 150 edges in this list (all with an increase in correlation greater than 0.26) was used to visualize the disease-specific coexpression network. We used Gephi version 0.9.2 (https://gephi.org/) to visualize the network^66^.

### Statistical analysis and implementation

We use the scientific Python toolkit, including the scipy and numpy Python packages^65^, to compute the statistical tests described in the manuscript, including Spearman correlation and associate *P*-values. *P*-values listed as less than 10^−308^ indicate values returned by the statistical software below the minimum nonzero floating-point value representable by the machine.

### Runtime and memory profiling

We used Python’s time module to obtain runtime measurements and used the top program in Linux (Ubuntu 17.04) to make periodic memory measurements. We made use of default scientific Python parallelism. We benchmarked our pipelines on a Google Cloud Enterprise instance with 16 logical cores and 104 gigabytes of memory and, for memory-inefficient alternative algorithms (**Supplementary Table 1**), on a local 2.30 GHz Intel Xeon E5-2650v3 with 48 logical cores and 384 GB of RAM. scVI was trained on a Nvidia Tesla V100-SXM2 with 16 GB of RAM.

## Data Availability

We used the following publicly available datasets:

- Notochord and neural plate cells from Cao *et al*.^1^ (GSE119945)
- Neurons from Mayer *et al*.^2^ (GSE104158)
- Neurons from Han *et al*.^3^ (https://figshare.com/articles/MCA_DGE_Data/5435866)
- Neurons from Zeisel *et al*.^4^ (http://mousebrain.org/)
- Neurons and microglia from Saunders *et al*.^5^ (GSE116470)
- In-situ hybridization images from the Allen Developing Mouse Brain Atlas^36^ (https://developingmouse.brain-map.org/)
- Bone marrow and cord blood cells from the Human Cell Atlas (https://preview.data.humancellatlas.org/)
- PBMCs from Zheng *et al*.^6^ (https://support.10xgenomics.com/single-cell-gene-expression/datasets)
- Fetal thymus hematopoietic cells from Zeng *et al*.^39^ (GSE133341)
- Microglia from Hammond *et al*.^8^ (GSE121654)
- Microglia from Masuda *et al*.^9^ (GSE124335)

## Acknowledgements

We thank R. Chun, B. DeMeo. C. Mak, S. Nyquist, C. Wong-Fannjiang, and the Berger and Bryson laboratory members for valuable discussions and feedback. B.H. is partially supported by NIH grant R01 GM081871 (to B. Berger) and by the Department of Defense (DoD) through the National Defense Science and Engineering Graduate Fellowship (NDSEG)

## Author Contributions

All authors conceived the algorithm. B.H. implemented the algorithm and performed the computational experiments. All authors interpreted the results and wrote the manuscript.

