## Supplementary Information for "Coexpression enables multi-study cellular trajectories of development and disease"

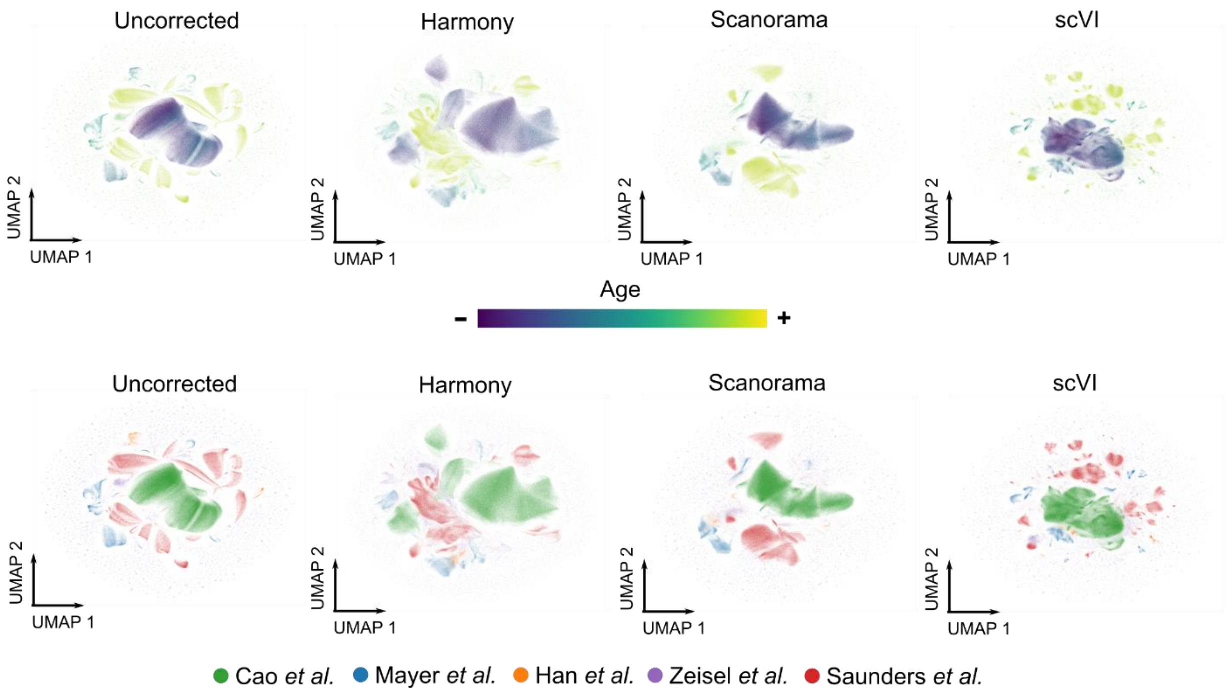


**Supplementary Figure 1: Neuronal cells in gene expression space across five studies do not organize according to development.**

Cells from five studies were concatenated and visualized in two dimensions using uniform manifold approximation and projection (UMAP). The visualization was learned from the log-transformed gene expression data ("Uncorrected"), Harmony-integrated embeddings ("Harmony"), Scanorama-corrected gene expression ("Scanorama"), and scVI-learned latent embeddings ("scVI") (**Methods**).

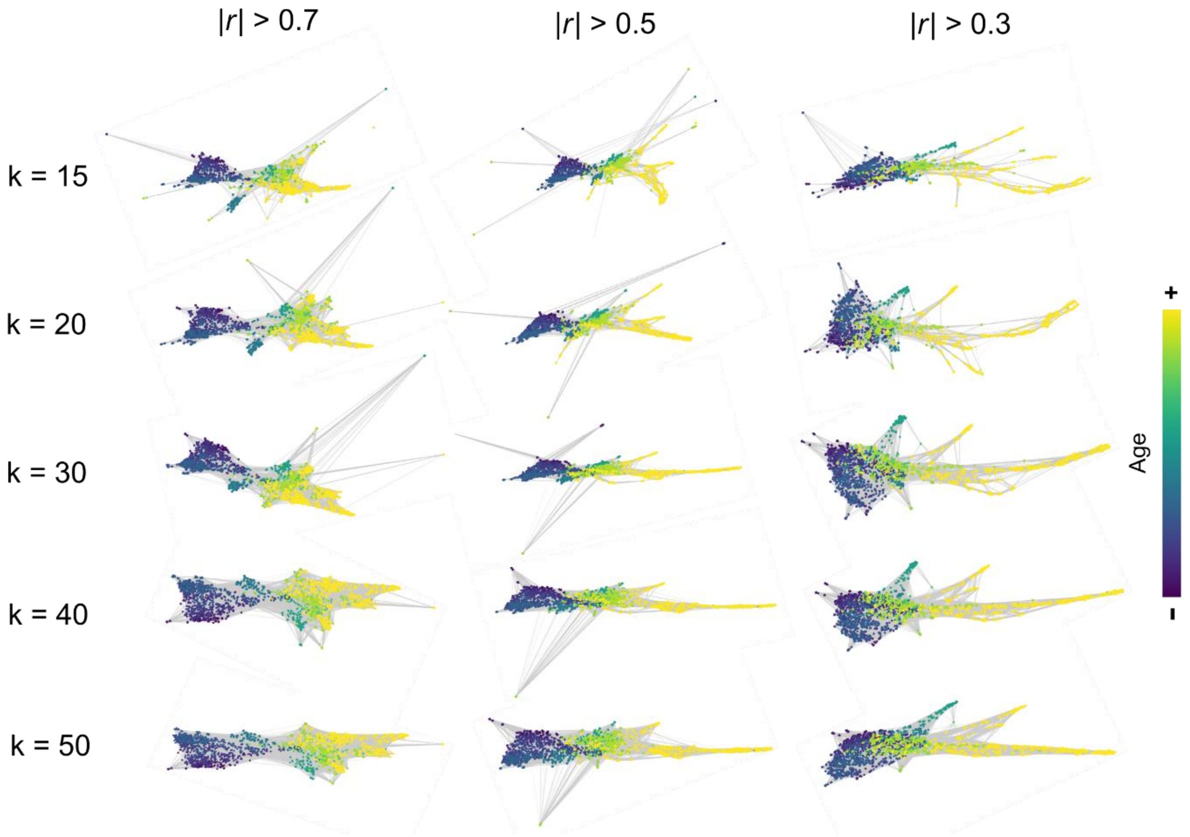


**Supplementary Figure 2: Coexpression landscape of mouse neuronal development while varying smoothing parameters.**

We visualized the coexpression landscape of the *k*-nearest neighbors graph at different values of *k* (vertical axis) and after different amounts of sparsification (horizontal axis). The networks were visualized in two dimensions using the ForceAtlas2 algorithm. Each point corresponds to a panresolution cluster and is colored by age using the same scheme as **Figure 2a**.

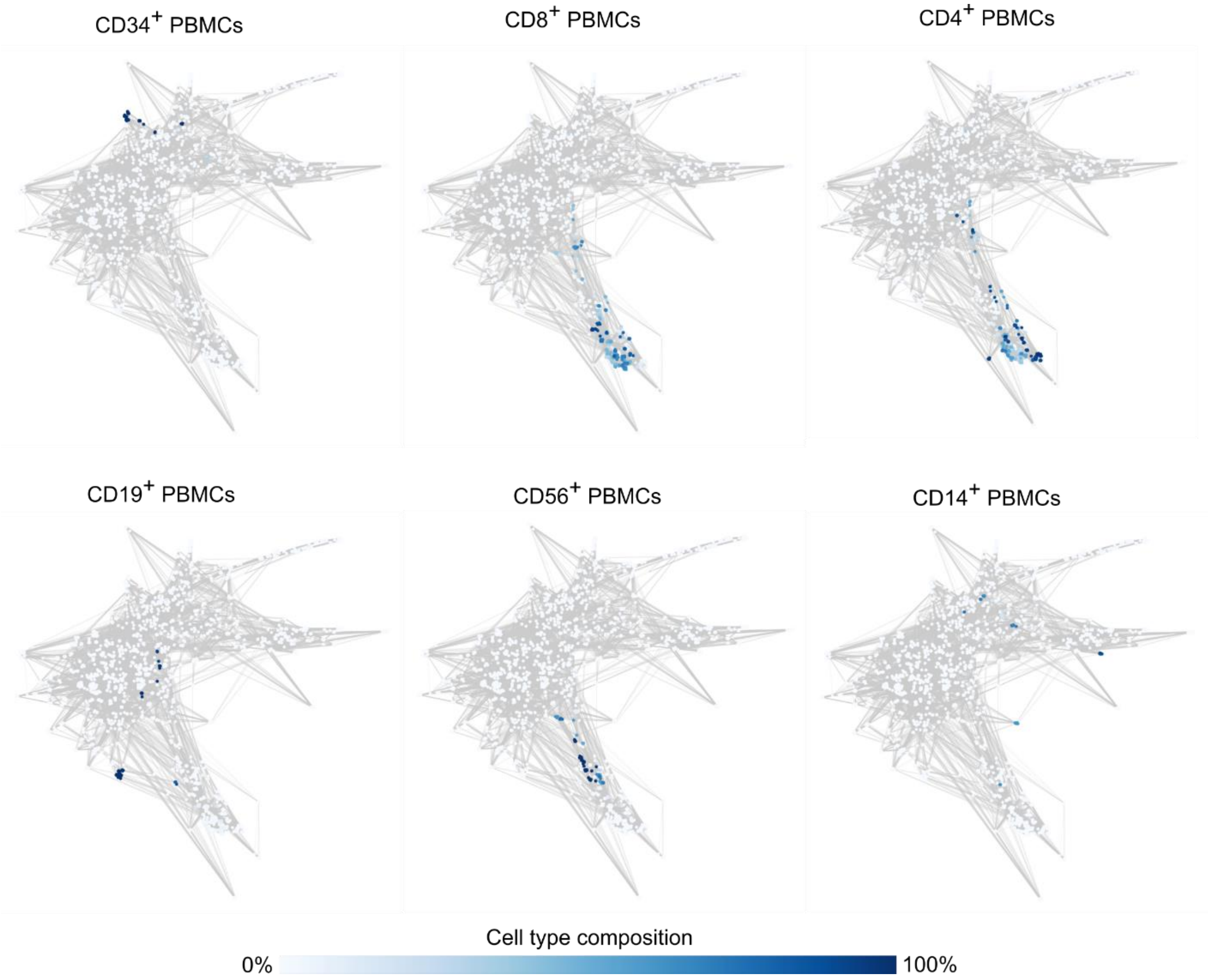


**Supplementary Figure 3: Proportion of FACS-labeled cells within panresolution clusters.** Panresolution clusters in the hematopoietic coexpression landscape were colored by their composition of FACS-labeled PBMCs, where labels were provided by the original study. Each coexpression landscape is plotted in the same coordinates as in **Figure 4**.

| **Method** | **Runtime (wall time)** | **Runtime (CPU time)** | **Peak memory usage** |
|---|---|---|---|
| Cutoff-based sparsification (this study) | 21 min | 168 min | 93.1 GB |
| No sparsification | < 21 min* | < 168 min* | > 378 GB$^{\dagger}$ |
| Lasso-based sparsification | > 24 hours$^{\ddagger}$ | > 24 hours$^{\ddagger}$ | 108.4 GB |

**Supplementary Table 1: Resource comparison between coexpression matrix computation strategies.**

Runtime (in both wall time and CPU time) and peak process memory usage for three different strategies for learning coexpression matrices across 2,442 panresolution clusters in our analysis of mouse neuronal development. "Cutoff-based sparsification" refers to the algorithm presented in this paper in which Pearson correlation matrices are learned for each panresolution cluster before setting correlations with an absolute value of less than 0.7 to zero and saving the result as a sparse matrix. "No sparsification" refers to the same procedure as above but without setting low associations to zero. This approach exceeds the memory of our benchmarking hardware and the process is killed before the full procedure is allowed to run. "Lasso-based sparsification" refers to learning finding a sparse matrix $\widehat{\mathbf{P}} = \underset{\mathbf{P}}{\operatorname{argmin}}\{\operatorname{tr}(\mathbf{\Sigma P}) - \log\det\mathbf{P} + \alpha||\mathbf{P}||_1\}$ where $\mathbf{\Sigma}$ is the sample covariance matrix and $\alpha$ is a parameter controlling the strength of the l1 regularization, otherwise known as the graphical lasso. Using the implementation from the sklearn Python package based on least angle regression for learning $\widehat{\mathbf{P}}$ for each panresolution cluster exceeds 24 hours of wall time. (*) The "no sparsification" procedure require fewer computational operations than applying an additional cutoff and therefore, assuming adequate memory resources, should theoretically require less overall time. (†) Procedure is killed upon exceeding 378 GB of memory. (‡) Process killed after 24 hours of wall time, extrapolated runtime is around 4 days.

| **Method** | **Runtime (wall time)** | **Runtime (CPU time)** | **Peak memory usage** |
|---|---|---|---|
| Loading, preprocessing, and joining datasets | 8 min | 10 min | 40.0 GB |
| Panresolution clustering | 27 min | 145 min | 41.9 GB |
| Coexpression matrix computation | 21 min | 168 min | 93.1 GB |
| Coexpression matrix dictionary learning | 6 min | 62 min | 9.9 GB |
| Constructing and analyzing coexpression landscape | 3 min | 3 min | 5.1 GB |
| Full pipeline | 65 min | 388 min | 93.1 GB |

**Supplementary Table 2: Resource usage of coexpression-based analysis of almost a million developing mouse neurons.**

Runtime (in both wall time and CPU time) and peak process memory usage for different stages of our analysis of mouse neuronal development. Additional details of the analysis are provided in **Methods**.
